# In Vitro Fertilization Accelerates Female Reproductive Aging Through Early Ovarian Failure

**DOI:** 10.1101/2025.10.13.682153

**Authors:** Eric A. Rhon-Calderon, Cassidy N. Hemphill, Alexandra J. Savage, Ana Domingo-Muelas, Zhengfeng Liu, Christopher J. Krapp, Laren Riesche, Nicolas D. Plachta, Richard M. Schultz, Marisa S. Bartolomei

## Abstract

Reproductive aging is characterized by the progressive decline of reproductive function, with broad implications for overall health and longevity. Environmental factors, including assisted reproductive technologies (ART), can accelerate reproductive aging by promoting premature ovarian failure in females. In vitro fertilization (IVF), though widely used and generally considered safe, is associated with lasting effects on offspring health. Using a mouse model that closely approximates human IVF, we demonstrate that IVF accelerates reproductive aging in female offspring by inducing premature ovarian failure. IVF-conceived females exhibit altered ovarian function, disrupted endocrine profiles, and transcriptomic and epigenetic changes consistent with premature reproductive decline. These findings reveal long-term consequences of IVF on female reproductive health and highlight the need to understand how early-life interventions influence reproductive longevity.

## Main

Reproductive aging is the physiological decline of biological functions essential for reproduction and overall health^1^. In females, reproductive aging involves the gradual decline in egg quality and quantity, accompanied by a decrease in reproductive hormones, e.g., estradiol, ultimately leading to menopause in humans or reproductive senescence in rodents^1^. In some women, reproductive aging occurs prematurely, manifesting as primary ovarian insufficiency (POI), a condition characterized by the loss of ovarian function before the age of 40^2^. POI represents an extreme manifestation of premature reproductive aging, which encompasses earlier and accelerated declines in ovarian reserve and endocrine function that negatively impact overall health and fertility, and are driven by genetic, environmental, and epigenetic factors^3^. Adverse conditions during early development, including those associated with assisted reproductive technologies (ART), could contribute to premature reproductive aging by disrupting epigenetic programming and metabolic processes^4–6^. These alterations can impair cellular function, increase mitochondrial stress, and lead to the early onset of age-related conditions, including reduced reproductive functions.

ART are non-coital methods of conception used to treat infertility^7^. Since the first successful human birth from *in vitro* fertilization (IVF) in 1978^8^, IVF and intracytoplasmic sperm injection have become the most widely used technologies to successfully achieve fertilization *in vitro*^8^. In both humans and mice, ART procedures are generally considered safe, but recent studies have shown that ART pregnancies are associated with higher risks of perinatal, neonatal, and placental adverse outcomes, rare genetic syndromes, and male offspring reproductive outcomes^9^.

IVF-associated adverse outcomes may be linked to critical developmental windows of gamete and preimplantation embryo epigenetic reprogramming, which are highly susceptible to external stresses^9,10^ and strongly influenced by the duration and conditions of embryo culture^11^. Despite ongoing refinements in IVF procedures to enhance fertilization and pregnancy rates, improvements in offspring outcomes have not been successful. The increasing use of IVF, driven by delayed childbearing, same-sex couples seeking parenthood, and rising infertility rates, underscores the need for further study on potential adverse outcomes for future generations^12^. Understanding the risk of premature reproductive aging in IVF-conceived individuals is crucial for evaluating long-term health risks and refining assisted reproductive practices.

In humans, female offspring conceived through IVF generally exhibit normal health, but recent research has identified adverse changes in cardiovascular and metabolic parameters^13^. Additionally, small cohort studies in adolescent females have reported a reduced ovarian reserve, altered timing of puberty, and hormonal imbalances^14^. Despite these findings, the overall impact of IVF on the female reproductive system remains unknown, and the mechanisms driving these adverse outcomes are yet to be fully elucidated.

Mice have become essential models for studying human disease and reproductive biology, and have been instrumental in the development and refinement of ART^15^. Mouse studies have shown that IVF is associated with increased risks of metabolic syndrome in offspring, including altered glucose, insulin, triglyceride, and cholesterol levels, as well as changes in liver transcriptomes and proteomes compared to naturally conceived offspring^4–6,16–21^. Furthermore, our lab has shown that IVF disrupts the reproductive system in male offspring, raising concerns about female reproductive effects^22^.

Despite observed adverse outcomes in multiple fetal and adult organs of IVF-conceived mice^4,16,23,24^, to date, no study has focused on the female reproductive system and the risk of premature aging. To address this gap, we examined the morphological and molecular effects of IVF on the ovaries and oocytes of female offspring and evaluated whether these alterations are associated with an increased risk of premature reproductive aging. Our findings reveal that IVF-conceived females display changes consistent with accelerated reproductive aging.

## Results

### Reduced estrogen levels and altered DNA methylation are associated with premature reproductive aging in IVF offspring

To investigate reproductive function in female IVF offspring, we examined ovaries using our mouse model (Fig. 1a). At 12 and 39 weeks, body and ovarian weights were significantly higher and lower, respectively, in IVF females compared to naturally conceived controls (Natural), resulting in a reduced ovarian-to-body weight ratio (Fig. 1b–d). The reduced ovarian weight could adversely affect ovarian function because ovaries are the primary producers of several key female sex hormones, including estradiol and progesterone^25^. Consistent with reduced ovarian weight, IVF females had lower serum estradiol levels at both 12 and 39 weeks and reduced progesterone at 12 weeks (Fig. 1e, f). Notably, estradiol and progesterone regulate follicular development and maintain an optimal microenvironment for oocyte growth^26^, and disruptions in estradiol levels can impair oocyte quality while changes in progesterone are associated with alterations in estrous cycle, uterine remodeling and future pregnancy complications^27^.

**Fig. 1:**
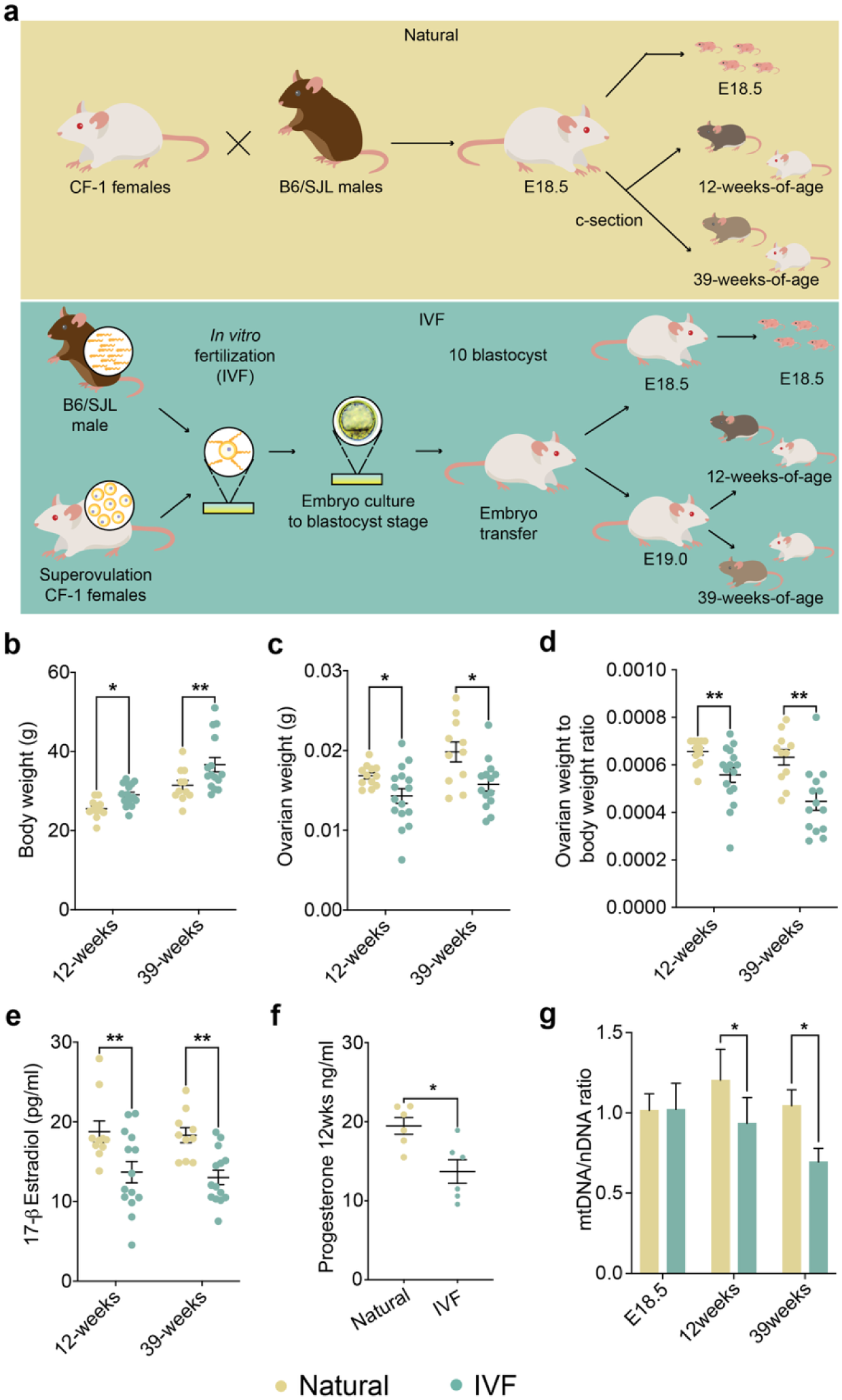
Mouse model and general parameters. **a,** Mouse model: Top: Naturally conceived embryos (Naturals) were generated using B6/SJL male mice and CF-1 female mice, and Bottom: IVF embryos were produced using capacitated sperm from B6/SJL mice and eggs from superovulated CF-1 mice. After embryo culture to the blastocyst stage, 10 blastocysts were transferred to pseudopregnant females. At E18.5, pregnant females from both groups were c-sectioned and fetuses were delivered and collected for molecular analysis. For adult cohorts c-section-delivered pups at E18.5 (Naturals) or E19.0 (IVF) were fostered with wild-type dams and monitored until 12 or 39-weeks of age. Shown are **b,** body weights before 6 h fasting; **c,** ovarian weights; **d,** ovary to body weight ratios; **e**, serum estradiol; and **f**, progesterone levels assayed by ELISA; **g,** The ratio of mitochondrial DNA relative to nuclear DNA assayed by RT-qPCR is depicted for all time points (see methods for primers). Data are depicted as mean±s.e.m, *n*=10-12 per group. The black line represents the mean of each group. Statistical significance was determined by t-test, \**P*<0.05, and \*\**P*<0.01 when IVF groups compared to Naturals.

Changes in mitochondrial function are considered a hallmark of aging^28^, where both mitochondrial DNA (mtDNA) copy number relative to nuclear DNA (ntDNA) and mitochondrial function decline with age in various tissues, including the ovaries^28^. To determine whether female IVF offspring exhibit mtDNA changes, we performed RT-qPCR using a mitochondrial DNA marker normalized to nuclear DNA. While no significant differences were observed at E18.5, we detected a significant decrease in the mtDNA/ntDNA ratio at both 12 and 39 weeks (Fig. 1g), suggesting the onset of an aging phenotype in IVF female offspring.

Collectively, these findings suggest that IVF contributes to premature ovarian aging by reducing estrogen levels and mtDNA content—each of which has been linked to ovarian dysfunction, mitochondrial impairment, and premature aging.

### Altered ovarian morphology is associated with premature reproductive aging in IVF offspring

Ovarian changes can impair oocyte maturation and subsequent ovulation^29^. Furthermore, early depletion of ovarian follicles is associated with reductions in ovarian size and estradiol production^30^. Thus, adverse effects on the ovarian reserve or ovarian function may compromise folliculogenesis, ultimately impacting female fertility. First, we quantified germ cells at E18.5 using whole-ovary staining for GCNA1 (TRA98), a germ cell marker^31^, and observed no difference between IVF and naturally conceived ovaries (Fig. 2a, b). In contrast, hematoxylin– eosin staining of whole ovaries revealed fewer primordial, primary, and secondary follicles at both 12 and 39 weeks (Fig. 2c–e). The absence of differences in germ cell number at E18.5, together with the reduction in follicles postnatally, suggests that accelerated germ cell depletion occurs after birth and may result from transcriptional dysregulation in somatic component of the ovary that is essential for normal follicle activation and development.

**Fig. 2:**
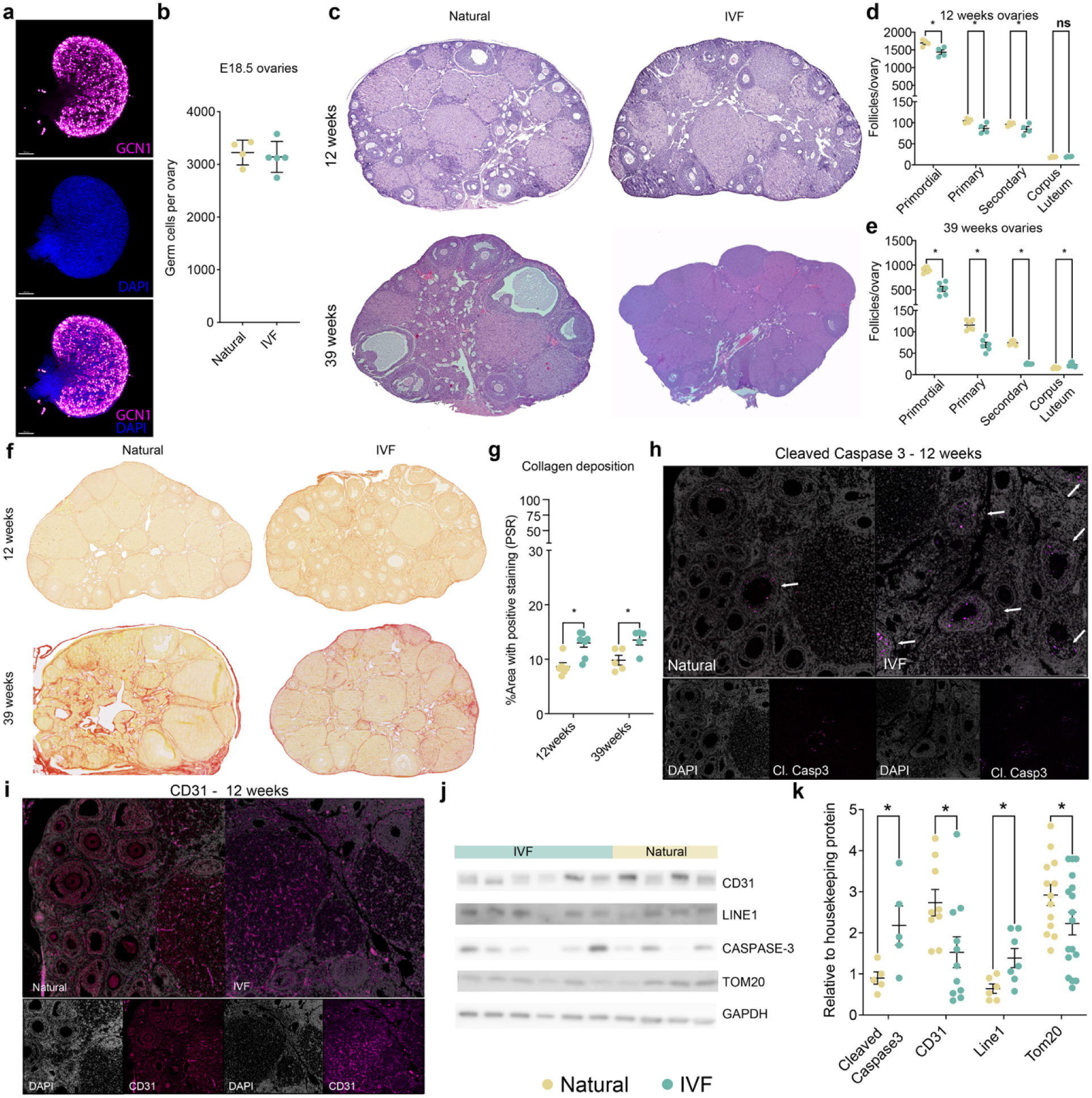
Ovarian morphology analysis. **a,** Whole ovarian staining at E18.5, GCNA1 for germ cells and DAPI for nuclei staining. **b,** Quantification of whole ovarian staining at E18.5. **c,** Ovarian cross-sections stained with hematoxylin-eosin at 12 and 39 weeks. **d,** Follicle number per ovary at 12 weeks. **e,** Follicle number per ovaries at 39 weeks. **f,** Ovarian cross-sections stained with picrosirius red (PSR) at 12 and 39 weeks. **g,** Percent of positively stained area with picrosirius red (PSR). **h,** Immunofluorescence for Cleaved Caspase 3 at 12 weeks. **i,** Immunofluorescence for CD31 at 12 weeks. **j,** Representative immunoblot. **k,** Quantification of immunoblot for Cleaved Caspase3, CD3, LINE1 and TOM20 at 12 weeks. Each data point represents an individual conceptus from a minimum of five different litters (n=5/group). Data are expressed as mean±s.e.m. Statistical analysis between groups was done by t-test and * represents a significant difference of *P*<0.05, when comparing IVF against Naturals.

Collagen is a key structural component of the ovary that supports follicle development and oocyte maturation, but excessive deposition with age leads to fibrosis and impaired ovarian function^32,33^. To assess whether IVF influences collagen accumulation, we performed picrosirius red staining and observed increased collagen density in IVF ovaries at both 12 and 39 weeks compared to controls (Fig. 2f, g). Ovarian aging and POI are also associated with increased cleaved caspase 3, which is a marker for increased atresia and apoptosis^34^ and reduced microvessel density^35^. We therefore stained ovaries with anti–cleaved caspase-3 and anti-CD31 antibodies to assess changes in apoptosis and endothelial cells, respectively, and observed increased caspase-3 and decreased CD31 by immunofluorescence (Fig. 2h, i). Additionally, immunoblotting revealed similar trends for cleaved caspase-3, CD31, and other aging markers, including LINE1 (Orf1p) and TOM20 (a mitochondrial marker) (Fig. 2j, k). Increased LINE1 activity is linked to elevated double-strand DNA breaks, mitochondrial dysfunction, and, consequently, increased apoptosis^36^. This increase may be a major driver of the mitochondrial dysfunction and apoptosis observed in our model, as evidenced by reduced mtDNA levels and elevated cleaved caspase-3 levels.

Taken together, these results indicate that IVF induces significant alterations in ovarian morphology, including a reduced primordial follicle pool, increased collagen deposition, heightened follicle atresia, and disrupted microvessel density in IVF offspring compared to naturally conceived controls, consistent with features of primary ovarian insufficiency.

### Ovarian Transcriptome in IVF Offspring Reveals Genes Linked to Premature Reproductive Aging

To determine whether the phenotypic changes described above were associated with altered gene expression and age-related patterns in IVF-conceived females, we performed RNAseq on whole ovaries collected at E18.5, 12, and 39 weeks. Principal component analysis (PCA) revealed clear clustering differences between IVF and naturally conceived groups (Extended Data Fig. 1a–c). To define age-dependent gene expression dynamics, we applied Within-Cluster Sum of Squares (WCSS) analysis and identified nine temporal clusters spanning E18.5 to 39 weeks (Fig. 3a). In naturally conceived females, these clusters followed predictable developmental trajectories, with subsets of genes decreasing (clusters 1, 6, 7, 9) or increasing (clusters 2, 3, 4, 5, 8) across age. By contrast, IVF females exhibited widespread deviations, with altered expression trajectories in all the clusters (Fig. 3a).

**Fig. 3:**
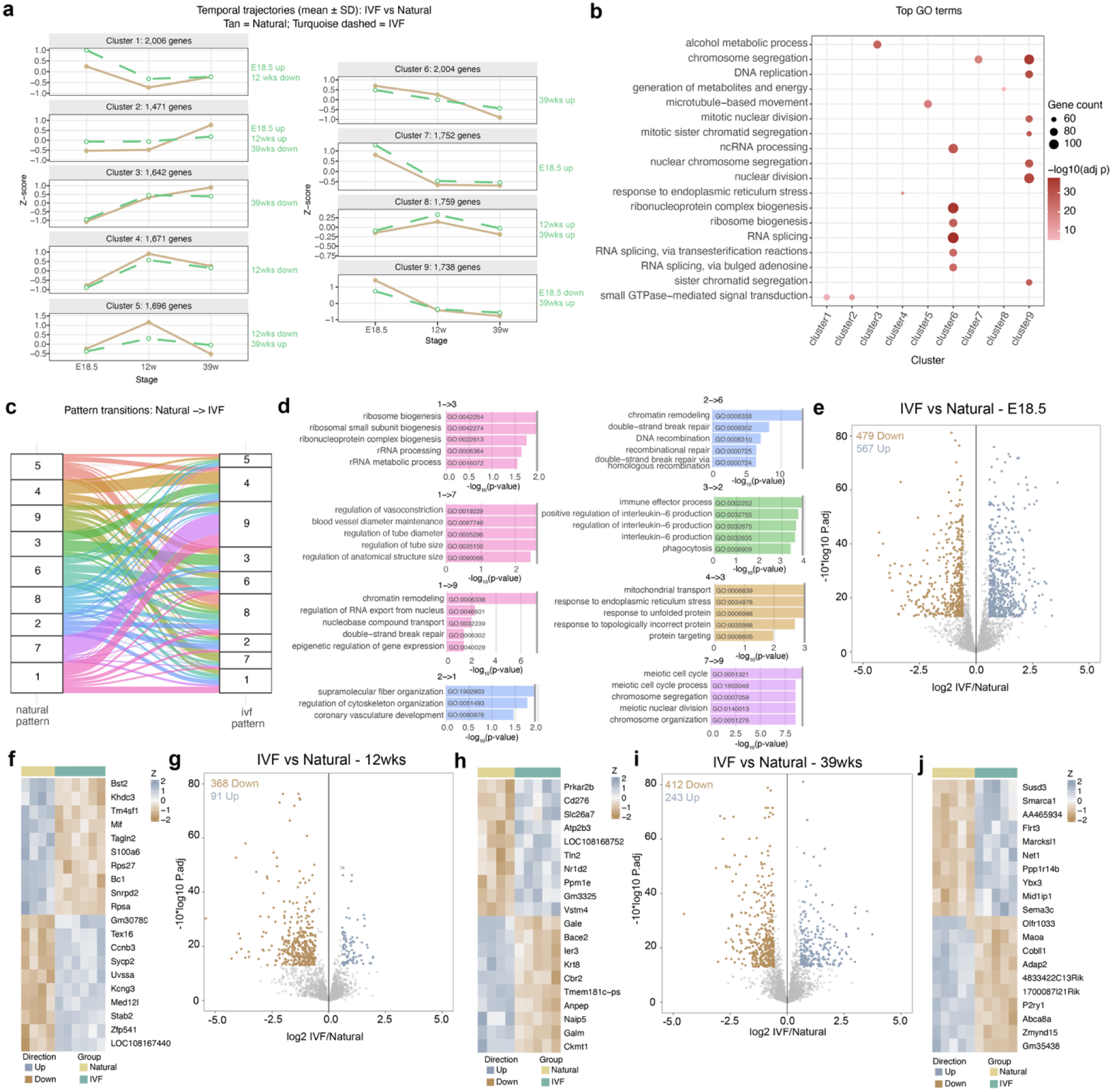
Transcriptome analysis of ovaries by RNAseq at three developmental timepoints. Within-Cluster Sum of Squares (WCSS) analysis using raw counts from RNAseq data at E18.5, 12 and 39 weeks: **a,** nine temporal expression patterns (k=9) across mouse ovarian development, using Natural as the training group. **b,** Top 20 enriched biological processes associated with ovarian function and their respective temporal expression cluster. **c,** Changes of temporal expression patterns for all genes from Natural to IVF group. **d,** Top 5 enriched biological process using the genes that change temporal expression patterns (clusters). Differential expression analysis using DeSeq2 for E18.5 gonads: **e,** Volcano plot. **f,** Heatmap with the most DEGs up or downregulated; 12-week ovaries: **g,** Volcano plot. **h,** Heatmap with the most DEGs up or downregulated; 39-week ovaries: **i,** Volcano plot. **j,** Heatmap with the most DEGs up or down. Color boxes at the top denote the experimental group.

To assign potential functionality to these clusters, Gene Ontology (GO) analysis was performed to determine whether genes within the nine clusters represented specific biological processes or pathways (Fig. 3b, Supplemental Table 1). Remarkably, several clusters showed strong associations with functions linked to ovarian aging and primary ovarian insufficiency (POI). For example, cluster 1 and 2 genes were enriched for signal transduction and homeostasis, indicating disruption of mechanisms essential for tissue homeostasis and aging^37^. Clusters 3, 4, and 5 downregulated at 39 weeks, were enriched for metabolic, autophagic, and mitochondrial pathways, consistent with impaired cellular energy regulation in aging ovaries. Clusters 7 and 9, increased and decreased at E18.5 respectively, were associated with replication, cell cycle, and chromosome segregation, reflecting early follicle loss and defective folliculogenesis, hallmarks of premature reproductive aging and POI^38^.

We further interrogated genes that shifted their affiliated cluster under IVF conditions (Fig. 3c). The alluvial plot demonstrated widespread reprogramming of temporal trajectories for pathways involved in metabolism, chromatid maintenance, chromosome integrity, cell cycle regulation, and respiration, processes that represent core hallmarks of reproductive aging^38^ and provide a mechanistic link to the follicular atresia, reduced oocyte quality, and diminished ovarian reserve observed in IVF offspring (Fig. 3d).

We next performed differential expression analysis and identified 567 upregulated and 479 downregulated genes at E18.5 (Fig. 3e; Supplemental Table 2), 91 upregulated and 368 downregulated genes at 12 weeks (Fig. 3g; Supplemental Table 3), and 243 upregulated and 412 downregulated genes at 39 weeks (Fig. 3i; Supplemental Table 4). The largest number of DEGs was observed at E18.5, and this difference did not appear to worsen with age. Several DEGs are directly linked to ovarian health and POI. *Hfm1* and *Spata22*, both required for homologous recombination and meiotic progression, were dysregulated, consistent with the impaired oocyte quality and follicle loss observed^39^. *Star*, which mediates cholesterol transport for estrogen biosynthesis, was altered in a manner consistent with the reduced estradiol levels detected in IVF females^40^. *Esr1*, encoding the estrogen receptor, was downregulated, potentially driving impaired follicle development and ovulation^41^. Additionally, *Gdf9*, a critical oocyte-derived growth factor for granulosa cell proliferation and folliculogenesis, was reduced, in line with the lower follicle counts observed^42^. Importantly, the dysregulation of genes involved in steroidogenesis and hormone signaling provides a molecular basis for the reduced estradiol and progesterone levels observed in IVF offspring. GO enrichment analysis of DEGs revealed pathways related to metabolism, mitochondrial function, cell cycle regulation, and gamete maturation at E18.5; metabolism and cell cycle regulation at 12 weeks; and metabolism and follicle development at 39 weeks (Extended Data Fig. 1e–g), highlighting progressive metabolic and developmental dysregulation across ovarian maturation stages. We next asked whether the identified DEGs were associated with established hallmarks of aging^43^. Pathway enrichment analysis revealed that genes dysregulated in IVF ovaries were significantly associated with multiple aging-related processes (Extended Data Fig. 1h,i). At 12 weeks, DEGs were enriched for pathways linked to altered intercellular communication, chronic inflammation, deregulated nutrient sensing, and genomic instability. By 39 weeks, these enrichments expanded to include mitochondrial dysfunction, loss of proteostasis, and stem cell exhaustion, hallmarks that together reflect progressive cellular decline.

We further identified the most upregulated and downregulated DEGs at each age (Fig. 3f,h,j), many of which play roles in ovarian homeostasis. Notably, intersection of the DEG datasets across all timepoints revealed two consistently altered genes, *Cdh13* and *Slc2a1* (Extended Data Fig. 1j,k), highlighting them as potential key mediators of IVF-associated ovarian dysfunction and accelerated reproductive aging. *Cdh13*, a cell adhesion molecule, has been associated with ovarian syndromes and may contribute to altered ovarian morphology^44^, while *Slc2a1* (Glut1), a glucose transporter essential for granulosa–oocyte energy transfer^45^, was consistently affected, providing a mechanistic link to the observed disrupted ovarian metabolism.

Taken together, these results align with the IVF-associated ovarian phenotypes and indicate that, as early as 12 weeks, the ovary exhibits transcriptional signatures of aging that intensify with time, suggesting that IVF accelerates molecular pathways driving reproductive aging.

### Single Oocyte Transcriptome in IVF Offspring Reveals Genes Linked to Premature Reproductive Aging

Given the cellular complexity of the ovary, we focused on germ cells using a modified SMART-Seq^46^ protocol that enabled sequencing of small numbers of primordial germ cells (PGCs), oocytes, or cumulus cells from E13.5, 12-week and 39-week females. As with whole ovaries, we first determined age-dependent gene expression dynamics in oocytes using WCSS analysis in both IVF and Natural groups. We identified nine clusters that captured gene expression trajectories from E13.5 to 39 weeks (Fig. 4a). In Natural females, clusters 1, 2, 5, 7 and 9 contained genes that decreased in expression from embryonic to adult stages; clusters 3, 4, 6, and 8 included genes that increased across development (Fig. 4a). By contrast, IVF females showed altered patterns in clusters 1, 2, 3, 4, 5 and 8 at one or more timepoints (Fig. 4a).

**Fig. 4:**
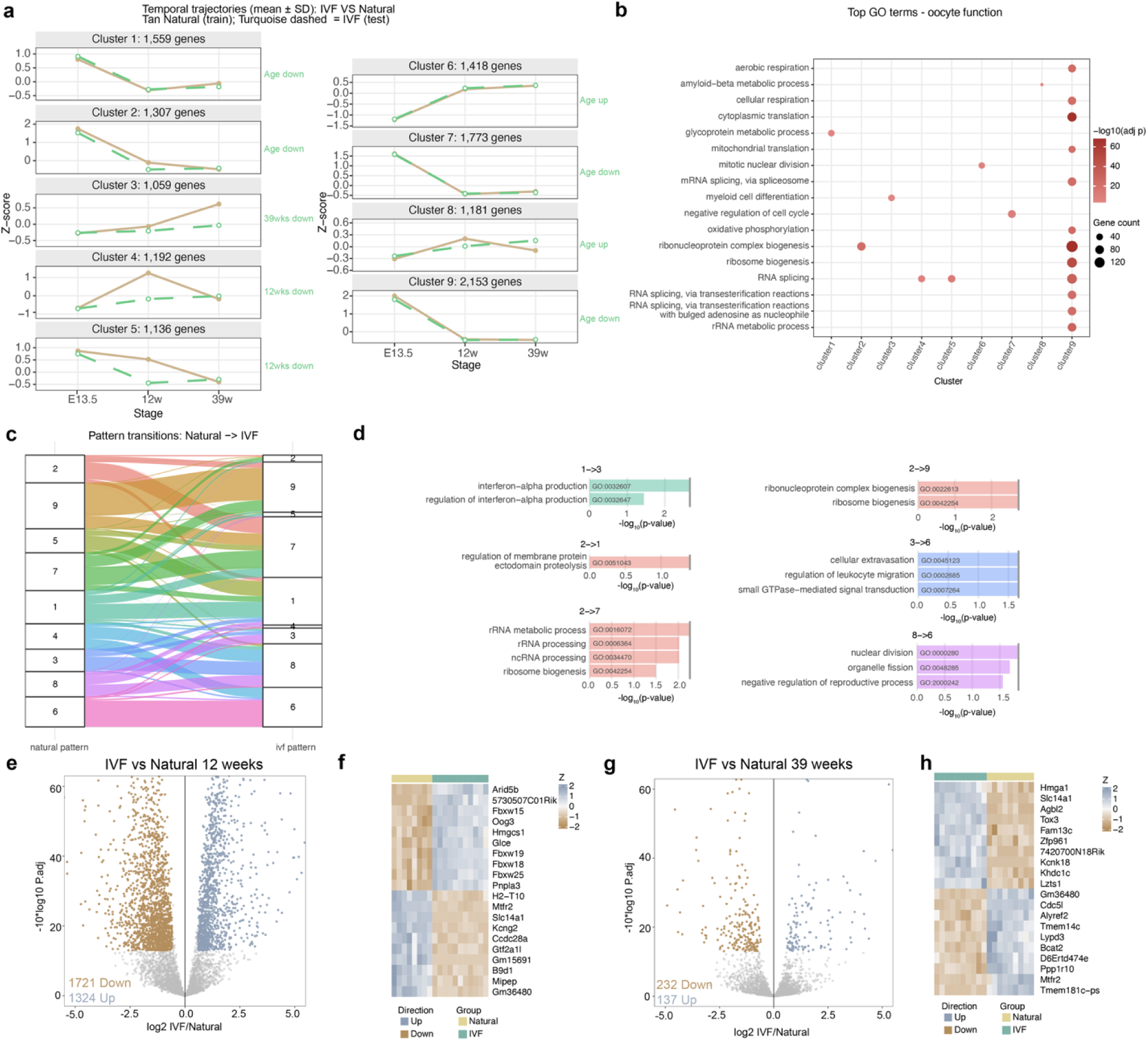
Transcriptome analysis of PGCs and single oocytes by RNAseq at three developmental timepoints. Within-Cluster Sum of Squares (WCSS) analysis using raw counts from RNAseq data at E13.5, 12 and 39 weeks: **a,** Temporal expression patterns (k=9) across oocyte development, using Natural as the training group. **b,** Top 20 enriched biological processes associated with oocyte function and their respective temporal expression cluster. **c,** Changes of temporal expression patterns for all genes from Natural to IVF groups. **d,** Top enriched biological process using the genes that change temporal expression patterns (clusters) that are associated with ovarian function. Differential expression analysis using DeSeq2 at E13.5 PGCs: **e,** Volcano plot. **f,** Heatmap with the most DEGs up or downregulated; 12-week oocytes: **g,** Volcano plot. **h,** Heatmap with the most DEGs up or down; 39-week oocytes: **i,** Volcano plot. **j,** Heatmap with the most DEGs up or downregulated. Color boxes at the top denote the experimental group.

As for whole ovaries, to assess the physiological relevance of these clusters, we performed GO analysis (Fig. 4b, Supplemental Table 5). Several clusters were strongly associated with biological pathways implicated in ovarian aging and primary ovarian insufficiency (POI). For example, cluster 2 genes, downregulated at 12 weeks, were enriched for ribosomal and translational pathways essential for oocyte maturation and fertilization. Ribosomal decline has been recognized as a hallmark of cellular and reproductive aging, reflecting impaired protein synthesis and reduced oocyte competence^47^. Cluster 5 genes, also decreased at 12 weeks, involved RNA modification and processing pathways, suggesting defects in post-transcriptional regulation that may compromise oocyte growth and follicle survival^48^. Cluster 8 genes, decreased at 12 and increased at 39 weeks in IVF, were enriched for metabolic and mitochondrial pathways critical for oocyte maintenance and energy homeostasis. Dysregulation of these pathways has been linked to oxidative stress, mitochondrial dysfunction, and premature ovarian failure^49^. Finally, Cluster 9 genes, enriched for stress response pathways, were downregulated, indicating diminished cellular resilience to metabolic and oxidative insults, a defining feature of aging oocytes and a key contributor to POI^47^. Together, these clusters highlight coordinated transcriptional disturbances in translation, RNA metabolism, energy regulation, and stress defense that collectively drive ovarian aging in IVF-conceived females.

As in whole ovaries, we analyzed genes that shifted their temporal expression patterns in the IVF group compared to Naturals (Fig. 4c). The alluvial plot illustrates how gene membership changed across the nine temporal clusters. GO analysis of the genes that switched patterns revealed enrichment for pathways involved in ribosomal transcription, notch pathways, chromatid maintenance, chromosome integrity, cell cycle regulation, and methylation, pathways that contribute to decreased oocyte quality^47^ (Fig. 4d).

Subsequent differential expression analysis revealed no DEGs at E13.5 (Extended Data Fig. 2a). Although PCA did not show clear separation between IVF and natural oocytes at 12 or 39 weeks (Extended Data Fig. 2b,c), we identified 1,721 upregulated and 1,324 downregulated genes at 12 weeks (Fig. 4e; Supplemental Table 6) and 232 upregulated and 137 downregulated genes at 39 weeks (Fig. 4g; Supplemental Table 7). The most upregulated and downregulated DEGs were identified, and heatmaps were generated to visualize their expression across the different groups (Fig. 4f,h).

GO enrichment analysis of DEGs revealed enrichment for pathways related to cell cycle regulation and metabolism at both 12 and 39 weeks (Extended Data Fig. 2e,f), highlighting disruptions in processes essential for oocyte quality, follicular maintenance, and overall ovarian function. We next examined whether dysregulated genes in oocytes were associated with hallmarks of aging (Extended Data Fig. 2g,h). At 12 weeks, IVF oocytes showed enrichment for pathways linked to cellular senescence, deregulated nutrient sensing through TOR signaling, genomic instability, loss of proteostasis, and mitochondrial dysfunction, indicating early activation of aging-associated stress responses. By 39 weeks, these transcriptional signatures intensified, with additional enrichment for chronic inflammation, mitochondrial respiratory defects, and stem cell exhaustion, consistent with progressive energetic decline and reduced oocyte quality.

Intersection of single-oocyte DEGs from 12 and 39 weeks revealed 22 overlapping upregulated genes in IVF (Extended Data Fig. 2i), including *Kif20a*, which is required for oocyte meiosis and early embryo development and is therefore essential for fertility^50^. The remaining upregulated genes were linked to folliculogenesis, steroidogenesis, the cell cycle, and other ovarian functions, although no direct correlation with POI has been described. In contrast, 49 downregulated genes (Extended Data Fig. 2j) included several with established roles in ovarian function and premature aging, such as *Lars2* (mutations cause Perrault syndrome with ovarian failure^51^), *Gpr3* (loss leads to premature ovarian aging and early meiotic resumption in mice^52^), *Lamc1* (associated with premature ovarian failure in human cohorts^53^), and *Drosha* (polymorphisms associated with idiopathic POI risk^54^). Together, these findings demonstrate that IVF induces persistent molecular signatures of aging in oocytes, characterized by disrupted mitochondrial metabolism and impaired proteostasis, which may underlie the accelerated reproductive decline observed in IVF-conceived females.

### Genes Linked to Premature Reproductive Aging are Detected in Transcriptomes from IVF Offspring Cumulus Cells

Cumulus cells, which provide critical support functions for oocyte maturation and reproduction, were analyzed using a SMART-Seq protocol at 12 and 39 weeks. In contrast to germ cells, PCA of cumulus cells (present only postnatally) showed clear separation between IVF and Natural groups (Extended Data Fig. 3e). Differential expression analysis revealed 204 upregulated and 256 downregulated genes at 12 weeks (Fig. 5a; Supplemental Table 8) and 57 upregulated and 676 downregulated genes at 39 weeks (Fig. 5b; Supplemental Table 9). The most upregulated and downregulated DEGs were depicted heatmaps and used for pathway enrichment trends (Fig. 5b, f). GO analysis revealed enrichment for pathways related to immune response, cell cycle, cell death, metabolism, and energy production. Among the DEGs, several genes previously associated with POI and reproductive aging in human granulosa cells were identified. These included *Inha*, linked to premature ovarian failure^55^; *Alkbh5*, whose downregulation is associated with defects in oocyte division and maturation^56^; and *Msi2*, whose reduced expression leads to decreased ovarian size and impaired folliculogenesis^57^, consistent with our observed phenotype. *Notch2* downregulation, associated with disrupted folliculogenesis and reduced fertility^58^, was also observed, as was *Egr1* upregulation, a known marker of ovarian aging^59^. Additionally, *Star* and *Cyp11a1*, key regulators of steroidogenesis, were dysregulated, supporting early endocrine disturbances associated with premature ovarian aging^40^. We then assessed whether transcriptional changes in cumulus cells reflected hallmarks of aging (Extended Data 3e,f). At 12 weeks, IVF-derived cumulus cells exhibited enrichment for pathways related to altered intercellular communication, cellular senescence, and chronic inflammation, indicating early disruption of oocyte–somatic signaling. By 39 weeks, these signatures broadened to include deregulated nutrient sensing, genomic instability, mitochondrial dysfunction, and stem cell exhaustion, consistent with cumulative cellular stress and loss of follicular support.

**Fig. 5:**
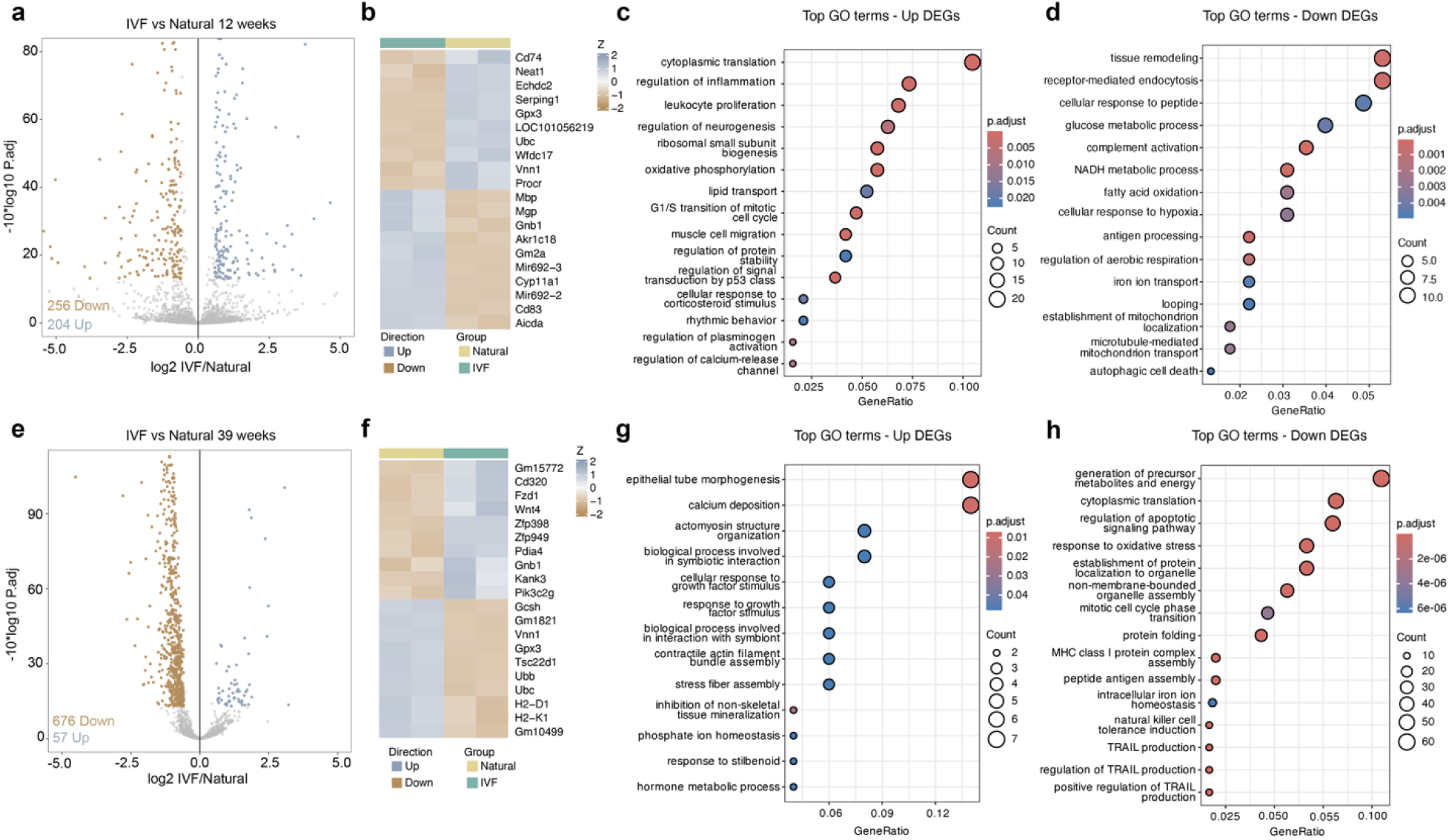
Transcriptome analysis of cumulus cells by RNAseq at two developmental timepoints. Differential expression analysis using DeSeq2 at 12 weeks: **a,** Volcano plot. **b,** Heatmap with the most DEGs up or down. **c,** Top gene ontology pathways using upregulated genes. **d,** Top gene ontology pathways using downregulated genes; 39-weeks: **e,** Volcano plot. **f,** Heatmap with the most DEGs up or down. **g,** Top gene ontology pathways using upregulated genes. **h,** Top gene ontology pathways using downregulated genes. Color boxes at the top denote the experimental group.

Finally, as before, we overlapped our datasets, revealing 11 upregulated and 84 downregulated DEGs common between 12- and 39-week cumulus cells (Extended Data Fig. 3g,h). Among these, two downregulated genes have previously been associated with premature ovarian insufficiency (POI): *Gja1*, which is essential for oocyte–cumulus communication and implicated in polycystic ovarian syndrome, and *Hif1a*, a regulator of follicular development via hypoxia pathways. Together, these data suggest that IVF imposes lasting transcriptional and metabolic reprogramming in cumulus cells, contributing to impaired follicle maintenance and the premature ovarian aging phenotype observed in IVF-conceived females.

### IVF Offspring Exhibit Altered Ovarian DNA Methylation

Because IVF is associated with changes in DNA methylation in multiple tissues^5,22^, and reproductive aging can be assessed through changes in DNA methylation, we interrogated global DNA methylation using a luminometric methylation assay (LUMA) on ovary DNA at E18.5, 12 and 39 weeks. We observed statistically significant decreased DNA methylation in ovaries from IVF-conceived female offspring compared to Naturals at all timepoints (Fig. 6a-c). Additionally, loss of DNA methylation at transposable elements (TEs) has previously been linked to oocyte aging and follicle depletion^60^. Accordingly, we measured DNA methylation changes in LINE1 and IAP elements, two major TEs in the mouse genome, using a targeted bisulfite sequencing assay. We observed a decrease in LINE1 DNA methylation in both 12- and 39-week-old IVF offspring compared to Naturals (Fig. 6a-c), with no changes detected in IAP (Fig. 6a-c).

**Fig. 6:**
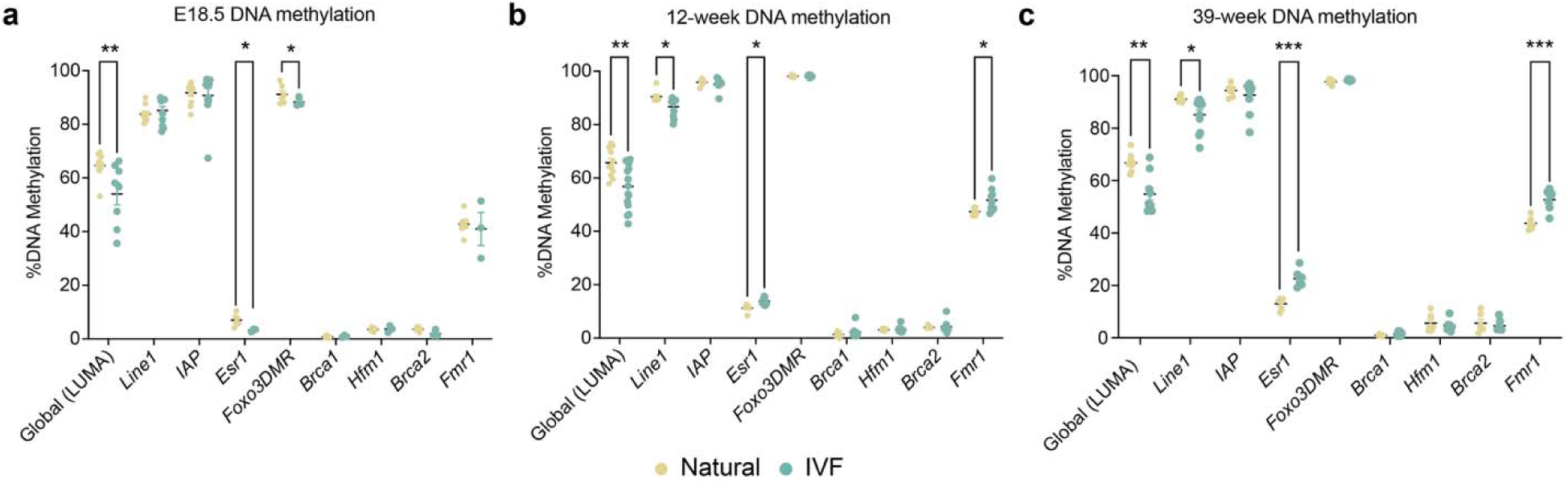
DNA methylation analysis of ovaries at three developmental timepoints. Global DNA methylation using luminometric methylation assay (LUMA) and whole ovarian DNA targeted bisulfite sequencing for specific loci (see methods section for CpG numbers) at for repetitive elements *Line1* and *IAP*, and promoter genes of *Esr1, Brca1, Hfm1, Brca2 and Fmr1,* and intergenic region of *Foxo3*: **a,** E18.5. **b,** 12 weeks. **c,** 39 weeks.

To determine whether gene expression changes were associated with altered DNA methylation, we performed targeted bisulfite sequencing at the promoters of genes linked to POI and differentially expressed across ages and sample types (*Esr1*, *Foxo3*, *Brca1*, *Hfm1*, *Brca2*, and *Fmr1*). DNA methylation at *Esr1* was significantly decreased at E18.5 (Fig. 6a), which may contribute to increased germ cell death during niche breakdown, and significantly increased at 12 and 39 weeks (Fig. 6b-c), correlating with downregulated expression and decreased serum estradiol levels. *Fmr1* methylation was also elevated at 12 and 39 weeks (Fig. 6b,c). Hypermethylation of *Esr1* and *Fmr1* is associated with menopause and senescence^61,62^, consistent with our phenotype. Because ovarian heterogeneity may mask cell type–specific transcription and DNA methylation changes, there may be an underestimation of observed differences. Taken together, these results suggest that DNA methylation dysregulation is a candidate for ovarian phenotypes, consistent with our previous findings in other tissues where altered DNA methylation mediates IVF-associated adverse outcomes^11,24^.

### IVF impacts the number of pups and resorptions in pregnancies from IVF female offspring

Because gene expression and ovarian health were disrupted in IVF-conceived females, we examined the fertility and pup ratio of in matings with IVF female offspring. Twelve-week-old IVF- and naturally conceived females were mated with CF-1 wild-type males, and concepti were either collected by c-sections at E18.5 (Fig. 7a) or allowed to naturally drop. Time to conception and gestation length (19 days) were comparable between groups. In contrast, pregnancies from IVF-conceived females showed a higher number of resorptions (Fig. 5b), fewer live pups 12 hours after birth (Fig. 7b), and reduced litter sizes across four consecutive litters (Fig. 7b). These results indicate that the changes observed in IVF female offspring have an impact on the number of pups and reabsorptions, likely due to decreased oocyte quality.

**Fig. 7:**
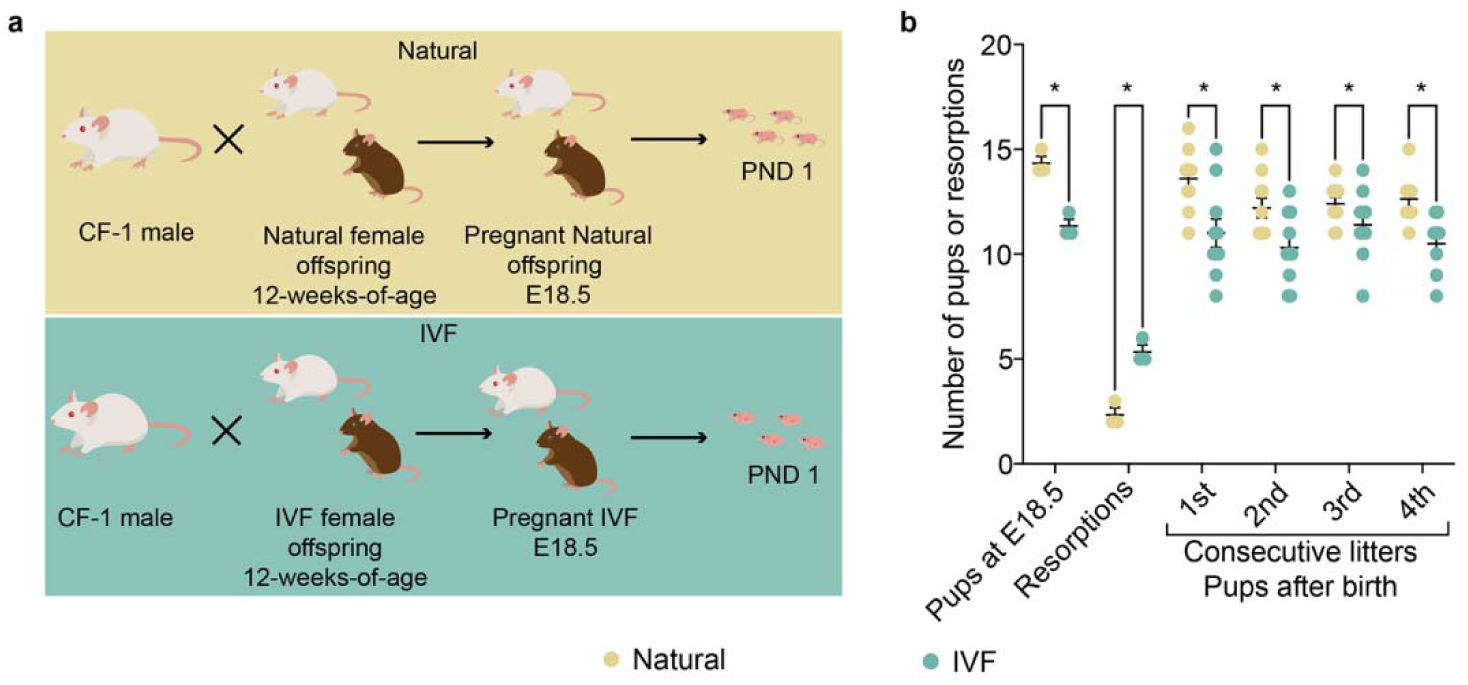
Number of pups and resorptions rates from IVF female offspring. **a,** Mouse model: Top: F2 Naturally conceived offspring were generated using CF-1 male and female offspring mice, and Bottom: F2 IVF offspring were produced by using CF-1 male and IVF female offspring mice. E18.5 pregnant females from both groups were c-sectioned and concepti were delivered. Other litters were allowed to naturally drop, and the number of pups were determined the day after delivery. **b,** Total number of pups is depicted for 4 consecutive litters, with data presented as summary of total pups after each litter from 4 breeding pairs/group.

## Discussion

In this study, we use a mouse model to demonstrate that IVF perturbs ovarian development and accelerates reproductive aging in female offspring, with changes in the number of pups and resorptions. These findings extend prior observations of ART-induced alterations in placental development, metabolism, and male reproductive health^4,6,24,63^, and provide new insight into how IVF impacts the female germline and associated cells across developmental stages.

A central theme emerging from our data is that IVF acts as a prenatal environmental exposure that reprograms ovarian biology. In humans, female IVF offspring appear to be at higher risk for bone aging abnormalities and exhibit changes in luteal hormone levels^14^. Here, we show that the hallmarks of premature reproductive aging, including reduced follicle counts, disrupted hormone production, and POI-associated gene expression, are evident in IVF females. Similar features are observed in human POI, as well as in models of maternal obesity and inflammation, supporting the idea that IVF mimics other adverse intrauterine environments in depleting the ovarian reserve^64^. Importantly, IVF-induced changes converge on pathways regulating metabolism, cell division, and folliculogenesis, raising the possibility that suboptimal conditions during early embryogenesis initiate a cascade that compromises ovarian function throughout life.

Our temporal clustering analyses revealed that IVF fundamentally alters age-dependent gene expression trajectories in ovaries and oocytes compared to naturally conceived females. While Natural groups showed predictable developmental transitions, such as declining expression of cell cycle and chromosomal pathways with age and stable expression of core metabolic regulators, IVF offspring exhibited significant deviations from these trajectories. Clusters that normally remained stable became dysregulated, with premature upregulation of DNA replication and cell cycle pathways in cumulus cells, and persistent activation of metabolic programs in both oocytes and somatic compartments. Conversely, clusters that typically increased with age, such as those supporting germ cell development and fertilization, were reduced in IVF, suggesting an early depletion of follicular support mechanisms. Together, these shifts reflect a reprogramming of ovarian developmental timing, with premature activation of stress and metabolic responses coupled to early loss of pathways required for long-term oocyte quality. Such remodeling of temporal expression dynamics provides a mechanistic framework for how IVF accelerates reproductive aging.

The cumulus cell findings indicate that IVF induces long-lasting transcriptional alterations that may underlie the premature reproductive aging observed in IVF offspring. As key regulators of oocyte maturation and follicular support, cumulus cells are particularly sensitive to disruptions in signaling and metabolism. The altered expression of genes such as *Inha*, *Alkbh5*, *Msi2*, and *Notch2* as early as 12 weeks, with the shared downregulation of *Gja1* and *Hif1a* at 12 and 39 weeks, points to impaired oocyte–cumulus communication and follicular homeostasis. Enrichment of immune, metabolic, and cell death pathways further suggests chronic cellular stress within the follicular microenvironment. Together, these changes provide a mechanistic link between IVF-induced molecular perturbations and the reduced follicle reserve, altered hormone levels, and accelerated ovarian aging seen in IVF-conceived females.

Our results also suggest that epigenetic dysregulation is one major mechanism linking IVF to premature reproductive aging. Altered DNA methylation at *Esr1* and *Fmr1*, together with global hypomethylation and dysregulation of LINE-1, are signatures of accelerated epigenetic aging^65^. Additionally, hypomethylation of LINE-1 elements is linked to its transcriptional activation, accumulation of DNA damage, and oocyte loss through apoptosis^66^. In humans, similar patterns are associated with ovarian insufficiency, menopause, and infertility^66^.

We observed changes in both the morphology and molecular status of the entire ovary and our analyses of oocytes and cumulus cells underscore the vulnerability of both germ and somatic ovarian compartments to IVF. Disruption of spindle formation, energy metabolism, and cumulus–oocyte communication pathways is similar to findings from obesity, alcohol, and diabetes exposure models, all of which compromise fertility^67–69^. The absence of differences in germ cell numbers at E18.5, followed by follicle loss later in life, indicates that epigenetic dysregulation within the somatic compartment of the ovary may impair chromatin remodeling and gene expression programs necessary for proper follicle activation, leading to accelerated postnatal germ cell depletion. This perspective has important implications for understanding how early-life perturbations can reprogram somatic–germ cell interactions and predispose to long-term ovarian dysfunction.

Taken together, our findings suggest a model in which IVF-induced stress during early embryogenesis disrupts female gonad development, with downstream effects on ovarian biology, at both germ and somatic cell levels. Notably, many of the pathways disrupted by IVF, mitochondrial function, oxidative stress responses, and senescence, are also central to biological aging^70^, raising the possibility that IVF accelerates systemic aging trajectories. Despite the increasing use of ART in humans, long-term studies on reproductive and metabolic health remain limited, especially in females. Our work highlights the utility of animal models for dissecting tissue-specific vulnerabilities and identifying candidate biomarkers for human studies. Understanding how IVF alters oocyte epigenetic integrity and ovarian somatic cell health will be essential for designing interventions that mitigate premature reproductive aging and improve fertility outcomes in ART populations. Ultimately, our findings underscore the importance of monitoring not only immediate pregnancy outcomes but also the lifelong health trajectories of ART-conceived individuals.

## Methods

### Animals

Breeding stocks of CF-1 females and CD-1 vasectomized males (from Charles River, USA) and B6/SJL males (from The Jackson Laboratory, USA), were maintained under pathogen-free conditions in polysulfone cages on a 12-hour light/dark cycle, with ad libitum access to water and chow (Laboratory Autoclavable Rodent Diet 5010, LabDiet).

### Generation of Natural offspring

Naturally conceived offspring (Natural) were obtained by mating 8-week-old CF-1 females, during their natural estrous cycle, with 8-week-old B6/SJL males. The presence of a vaginal plug was recorded as embryonic day 0.5 (E0.5), and embryos were allowed to develop entirely in vivo without embryo transfer (Fig. 1a).

### Generation of in vitro fertilization offspring

As previously described^24^, IVF offspring were generated according to optimized protocols recommended by the Jackson Laboratory^71^. Eight-week-old CF-1 females were superovulated with 5 IU eCG, followed 46 h later by 5 IU hCG. On the day of IVF, sperm were collected from the vas deferens and cauda epididymis of B6/SJL males into EmbryoMax Human Tubal Fluid medium (HTF, EMD Millipore) supplemented with 3% w/v bovine serum albumin (BSA; AlbuMax, GIBCO) and maintained under mineral oil (Irvine Scientific, USA). Sperm were capacitated for at least 1 h prior to fertilization. Oocytes were harvested and inseminated with capacitated sperm in HTF medium. After 4 h, fertilized oocytes with visible pronuclei were washed through HTF and then into EmbryoMax KSOM medium supplemented with half-strength amino acids (KSOM+AA, EMD Millipore), before being cultured to the blastocyst stage under mineral oil at 37 °C in 5% CO, 5% O, and 90% N. After 3.5 days of culture, blastocysts were washed in Multipurpose Handling Medium Complete (MHM-C, Irvine Scientific) containing Gentamicin prior to embryo transfer. Pseudopregnant recipients (3.5 days post-copulation) were generated by mating CF-1 females with CD-1 vasectomized males, and each recipient received ten blastocysts by non-surgical embryo transfer (Fig. 1b). The day of blastocyst transfer was designated as E3.5.

### Tissue collection

At E18.5 and at 12 or 39 weeks of age, naturally mated females or pseudopregnant recipients with IVF embryos were euthanized, and one ovary was snap-frozen in liquid nitrogen while the other was fixed in 10% phosphate-buffered formalin for histological analysis. Blood was also collected, and serum was isolated by centrifugation at 4 °C and stored at –80 °C for hormone measurements.

### Whole embryonic ovary staining

Fixed ovaries at E18.5 were stained for germ cell counting using a previously described protocol^31^. Briefly, after 24 hours of fixation, gonads were washed in fresh PBS for 30 minutes at room temperature on a lab shaker (100 RPM). Ovaries were then incubated in blocking solution (2% BSA, 0.5% Triton X-100, and 15% goat serum in PBS) for 3 hours at room temperature with shaking. Following blocking, ovaries were incubated for 4 days with anti-GCNA1 (1:100, Abcam, USA) diluted in blocking solution, under the same conditions. After primary antibody incubation, ovaries were washed four times for 15 minutes in blocking solution without goat serum to remove excess antibody. Samples were then incubated for 2 days with donkey anti-rat IgG (H+L) highly cross-adsorbed secondary antibody conjugated to Alexa Fluor™ 555 (1:1000, ThermoFisher, USA) in blocking solution. Ovaries were washed three times for 30 minutes in blocking solution without goat serum, followed by incubation with DAPI (1:100, ThermoFisher, USA) for 30 minutes in blocking solution. Finally, ovaries were washed in PBS and cleared using Scale CUBIC-1 solution (125 g urea [CAS 57-13-6], 156 g Quadrol 80% [CAS 102-60-3], 144 g water, followed by 75 g Triton X-100 [CAS 9036-19-5] after dissolution). Imaging was performed on a Leica SP8 confocal microscope, acquiring 3D scans of whole ovaries with a z-step size of 5 µm over 30 minutes. Germ cell quantification from three-dimensional ovarian visualizations was performed using Imaris 9.7 software (Bitplane AG). All analyses were conducted in a blinded manner, with samples coded and evaluated by an independent reviewer to prevent bias.

### Hematoxylin-eosin and Picrosirius red staining

Ovarian sections were stained with hematoxylin-eosin as previously described^72^. Stain sections were imaged using an EVOS FL Auto Cell Imaging System and software (Thermo Fisher Scientific, Waltham, MA, USA) at 4× magnification. Follicular counting was performed as previously described^72^ by two blinded people. To avoid multiple counting, every 10 sections, only follicles with clear oocyte nuclei were counted. Follicles were expressed by total number per ovary.

To assess changes in collagen deposition and potential fibrosis, picrosirius red (PR) staining was used on ovarian sections as previously described^22^. Briefly, tissue sections were deparaffinized in xylene and rehydrated through a graded series of ethanol, then immersed in a PR staining solution (Sirius Red F3B Direct Red 80, C.I. 35782, Sigma-Aldrich, St. Louis, MO, USA in a 1.3% saturated aqueous picric acid solution, Sigma-Aldrich, St. Louis, MO, USA). After a 30-min incubation in the staining solution at room temperature, the slides were destained with 0.05 N HCl. The tissue sections were then dehydrated in 100% ethanol, cleared in xylene, and mounted with Permount mounting medium. Stained slides were imaged using an EVOS FL Auto Cell Imaging System (Thermo Fisher Scientific, Waltham, MA, USA). The percentage of PR-positive signal was calculated using three different testicular sections per animal with ImageJ.

### Ovarian immunohistochemistry

For ovarian immunochemistry, ovaries were treated as previously published^24^. Briefly, histological sections were rehydrated using xylene, followed by graded series of ethanol and then with PBS. Antigen retrieval was performed using Antigen Unmasking Solution (Vector Laboratories, CA, USA). Sections were subsequently washed, quenched with a 30% hydrogen peroxide/methanol solution, washed again in PBS, and blocked using 15% normal goat serum in 0.3% Triton-PBS for 1 h at room temperature. The primary antibody, anti-CD31 (Abcam, MA, USA), anti-cleaved caspase 3 (Cell Signaling, USA), was applied at a dilution of 1:100 for 1 h at room temperature, followed by overnight incubation at 4°C. Negative control slides received 15% normal goat serum without primary antibody. The following day, slides were washed with PBS and secondary antibody was applied: Goat anti-rabbit IgG (H+L) Highly Cross-Adsorbed Secondary Antibody, Alexa Fluor™ Plus 647 (ThermoFisher, USA) at 1:250 for 1 h. After washing with PBS, slides were incubated with 4’,6-diamidino-2-phenylindole (DAPI, ThermoFisher, USA) at 1:100 in blocking solution for 30 min. Finally, slides were washed in PBS and distilled water, then mounted using ProLong™ Diamond Antifade Mountant (ThermoFisher, USA).

### Estradiol and Progesterone levels

Whole blood was collected by cardiac puncture on the day of euthanasia. Samples were centrifuged, serum was collected and stored a -80°C. Estradiol was measured using Mouse Estradiol ELISA kit (Abcam, USA) and progesterone was measured using a Mouse Progesterone ELISA kit (Crystal Chem, USA) following manufacturer instructions.

### DNA and RNA isolation from ovaries

DNA and RNA were isolated from one-half of each snap-frozen ovaries or a whole E18.5 gonad, as previously described^4^. Briefly, for DNA tissues were digested in lysis buffer (50 mM Tris, pH 8.0, 100 mM EDTA, 0.5% SDS) with proteinase K (180 U/mL; Sigma-Aldrich) overnight at 55°C. Genomic DNA was isolated using phenol:chloroform:alcohol (25:24:1; Sigma-Aldrich), followed by ethanol precipitation and resuspension in TE buffer (10 mM Tris-HCl, pH 8.0, 0.5 mM EDTA).

RNA was isolated using an adapted protocol of trizol (Thermo Fisher Scientific) and Qiagen Micro RNA Kit (Qiagen, Germany), following the manufacturer’s protocol. DNAse treatment was performed during RNA isolation to eliminate genomic DNA contamination. RNA quality and concentration were determined by RNA ScreenTape analysis using a TapeStation (Agilent Technologies, USA).

### Luminometric methylation assay

Genomic DNA (1 µg) from ovaries was used to measure global DNA methylation by luminometric methylation assay as previously described^73^.

### Locus specific DNA methylation analysis using targeted next-generation bisulfite-sequencing

DNA methylation levels at *Esr1*, *Foxo3*, *Brca1*, *Hfm1*, *Brca2*, *Fmr1,* LINE1 and IAP genomic regions were assessed using targeted DNA methylation analysis via next-generation sequencing. Primer sequences are provided in Supplemental Table 14. The assay was performed as previously described^74^.

### Immunoblotting

Half of the ovaries from 12- and 39-week females were used for protein analysis by Western Blot as previously described^4,24^. Briefly, ovaries were lysed in RIPA buffer (Cell Signaling, USA) containing protease inhibitors without EDTA (Merck Millipore, USA). Protein lysates (20 μg per sample) were separated and probed with primary antibodies diluted in 5% nonfat dry milk in TBS-T: anti-Tubulin (1:1,000; Cell Signaling, USA), anti-CD31 (1:100; Abcam, MA, USA), anti-Cleaved Caspase-3 (1:200; Cell Signaling, USA), anti-Orf1p-LINE1 (1:500; Abcam, Waltham, MA, USA), and anti-TOM20 (1:1,000; Abcam, USA). Levels of CD31, Cleaved Caspase-3, Orf1p-LINE1, and TOM20 were quantified relative to GAPDH (1:10,000; Cell Signaling, USA) and compared between groups.

### Mitochondrial DNA (mtDNA) copy number

Ovary mitochondrial DNA (mtDNA) copy number was estimated using a real-time quantitative polymerase chain reaction (RT-qPCR) as previously described^75^. Briefly, previously extracted DNA was amplified using primers that target two mitochondrially-encoded genes, ND1 and *Cox3* (Supplementary Table 15), and the nuclear small subunit 18S rRNA (18S). RT-qPCRs were performed using the 7 Flex Real-Time PCR System (Life Technologies) using the Power SYBR green master mix (Applied Biosystems, Foster City, CA, USA), with the optimized cycling parameters detailed in Supplementary Table 15. The mtDNA/nuclear DNA (nDNA) ratio was calculated using the ΔΔCt method.

### RNA sequencing

We performed RNA sequencing on a random subset of E18.5, 12 and 39-week ovaries from n=5/6 individuals for each group and different litters to determine changes in gene expression. As previously described^4,16,22^, total RNA (4 µg) was used to prepare mRNA-sequencing libraries using KAPA mRNA-Seq library synthesis kit and KAPA Single-Indexed adapter kit (Kapa Biosystems, Wilmington, MA, USA). Library quality control was conducted using High Sensitivity DNA ScreenTape for TapeStation (Agilent Technologies, USA) and Kapa Library Quantification Kit (Kapa Biosystems, Wilmington, MA, USA). Sequencing was performed using the NovaSeq 1000 platform (Illumina, San Diego, CA).

RNA-seq reads were analyzed as previously described^4^. The raw sequencing data reported in this work has been deposited in the NCBI Gene Expression Omnibus under accession number GSE307881. Count data were analyzed in R (version 4.4.1) using a combination of Bioconductor and CRAN packages. Differential expression analysis was performed with DESeq2, while temporal clustering of gene expression trajectories was assessed using a Within-Cluster Sum of Squares (WCSS) approach, as previously described^76^. Briefly, for clustering we first applied a stringent expression filter to the raw count matrix: genes were retained only if they reached CPM ≥ 1 (computed with edgeR::cpm) in ≥20% of all samples (minimum 2 samples) and were detected in ≥1 sample within each age group. Counts passing this filter were converted to CPM and transformed to gene-wise Z-scores (mean-centered and variance-scaled per gene). For each gene, we then computed stage means (mean Z per age) to obtain a genes-by-age profile matrix used for clustering. To choose the number of clusters, we ran k-means (R kmeans, nstart = 50, iter.max = 1000, fixed seed) across k = 2–15 and recorded the within-cluster sum of squares for each k. The optimal k was selected by the elbow criterion, i.e., the smallest k beyond which additional clusters yielded diminishing reductions in WCSS; as a quantitative aid, we also inspected the relative improvement and favored the first k with <5% gain. Functional enrichment of differentially expressed genes was carried out through gene ontology (GO) overrepresentation analysis. Changes in temporal cluster membership between groups were visualized with alluvial plots, and overlaps across datasets were examined using the VennDiagram package.

### Low input RNA sequencing for single oocytes and cumulus cells

We performed RNA sequencing on a subset of primordial germ cells (PGCs), single oocytes and cumulus cells from E13.5, 12- and 39-week-old mice. PGCs were collected as previously described^74^. For oocytes, seventy-two hours before their 12- or 39-week birthdate, IVF and naturally conceived female offspring were superovulated with 5 IU eCG, followed by 5 IU hCG 46 hours later. The next day, cumulus-oocyte complexes (COCs) were collected from the superovulated animals and deposited in MHM-C under mineral oil containing 10 µg/ml hyaluronidase (Sigma Aldrich, Germany) and incubated at 37°C in an atmosphere of 5% CO, 5% O, and 90% N for two minutes. Then, the cleaned oocytes were washed three times in fresh MHM-C drops and individually snap-frozen in 2 µl of TCL lysis buffer (2× TCL, Qiagen) with 1% 2-mercaptoethanol (Bio-Rad) in low-binding PCR tubes. A similar procedure was followed for cumulus cells; except they were counted using a hemocytometer to ensure 100 cells per 2 µl of solution before freezing. RNA sequencing libraries were constructed following a modified SMART-Seq protocol^46^. Briefly, after cell lysis, RNA was isolated using RNAClean-XP beads (Beckman Coulter, USA), and full-length polyadenylated RNA was reverse-transcribed using Superscript II (Invitrogen, USA). The cDNA was amplified using 10 cycles and subsequently (0.33 ng) used to construct a pool of uniquely indexed samples with the Nextera XT kit (Illumina, USA). Library quality control was conducted using High Sensitivity DNA ScreenTape for TapeStation (Agilent Technologies) and a Library Quantification Kit (KAPA Biosystems). Finally, pooled libraries were sequenced on a NextSeq 1000. Data was analyzed as shown before^46^. Briefly, raw reads were trimmed with trimgalore and aligned to the mm10 reference *Mus musculus* genome using Star. A count table was generated using featureCounts and used as input for the R package DESeq2. Finally, pathway analyses were performed using the clusterProfiler R package.

### Statistical analyses

All samples were statistically analyzed by t-test. Probability of genomic regions in the array were compared using a Bernoulli distribution using R v 4.2.2 (R foundation for Statistical Computing; www.R-project.org/). Significant differences comparing Natural and IVF offspring were denoted as statistically significant if *P*<0.05. Differences in variability was calculated by F-test, statistically significant differences comparing Natural and IVF were denoted if *P*<0.05. Sperm correlation analysis was performed using a Pearson r test and consider significant if *P*<0.05. All statistical analyses were performed using GraphPad Prism version 10.

### Study approval

All animal work was conducted with the approval of the Institutional Animal Care and Use Committee (IACUC) at the University of Pennsylvania. IACUC protocol 80354 has been previously revised and approved.

### Data availability

All values for all data points in graphs are reported in the Supporting Data Values file. Additionally, the data array and sequencing data underlying this article is available at NCBI Gene Expression Omnibus at https://www.ncbi.nlm.nih.gov/geo/, under GEO accession number GSE307881.

## Author Contributions

EAR-C and MSB designed research studies. EAR-C, CNH, AJS, ADM, CJK, ZL, and LN conducted experiments. EAR-C, AJS, ADM and CJK acquired data. EAR-C wrote the manuscript; and all authors edited and reviewed the manuscript prior to submission. EAR-C is listed as first author based on his conceptualization and initiation of the project.

## Acknowledgements

We acknowledge the individuals that helped to make this work possible, including Joanne Thorvaldsen, for her technical expertise; Ken Zaret for use of microtome; CHOP Pathology Core Laboratory for use of embedding equipment, Penn Cell and Developmental Biology Microscopy core for use of the imaging facilities. This work was funded by a National Centers for Translational Research in Reproduction and Infertility grant HD068157 (MSB) and HD102013 (NP), Ruth L. Kirschstein National Service Award Individual Postdoctoral Fellowship HD107914 (ER-C), Ruth L. Kirschstein National Service Award Individual Doctoral Fellowship HD117524 (CNH), and The National Institute of General Medical Sciences GM139970 (NP).

## References

1. Llarena, N. & Hine, C. Reproductive Longevity and Aging: Geroscience Approaches to Maintain Long-Term Ovarian Fitness. The Journals of Gerontology: Series A 76, 1551–1560 (2021).

2. Chon, S. J., Umair, Z. & Yoon, M.-S. Premature Ovarian Insufficiency: Past, Present, and Future. Front Cell Dev Biol 9, 672890 (2021).

3. Yureneva, S. et al. Searching for female reproductive aging and longevity biomarkers. Aging (Albany NY) 13, 16873–16894 (2021).

4. Narapareddy, L. et al. Sex-specific effects of in vitro fertilization on adult metabolic outcomes and hepatic transcriptome and proteome in mouse. FASEB J 35, e21523 (2021).

5. Lira-Albarrán, S., Liu, X., Lee, S. H. & Rinaudo, P. DNA methylation profile of liver of mice conceived by in vitro fertilization. J Dev Orig Health Dis 13, 358–366 (2022).

6. Donjacour, A., Liu, X., Lin, W., Simbulan, R. & Rinaudo, P. F. In Vitro Fertilization Affects Growth and Glucose Metabolism in a Sex-Specific Manner in an Outbred Mouse Model1. Biology of Reproduction 90, 80, 1–10 (2014).

7. Choe, J. & Shanks, A. L. In Vitro Fertilization. in StatPearls (StatPearls Publishing, Treasure Island (FL), 2024).

8. Eskew, A. M. & Jungheim, E. S. A History of Developments to Improve in vitro Fertilization. Mo Med 114, 156–159 (2017).

9. Rhon-Calderon, E. A., Vrooman, L. A., Riesche, L. & Bartolomei, M. S. The effects of Assisted Reproductive Technologies on genomic imprinting in the placenta. Placenta 84, 37–43 (2019).

10. Gosden, R., Trasler, J., Lucifero, D. & Faddy, M. Rare congenital disorders, imprinted genes, and assisted reproductive technology. The Lancet 361, 1975–1977 (2003).

11. Vrooman, L. A. et al. Placental Abnormalities are Associated With Specific Windows of Embryo Culture in a Mouse Model. Frontiers in Cell and Developmental Biology 10, (2022).

12. Kushnir, V. A., Smith, G. D. & Adashi, E. Y. The Future of IVF: The New Normal in Human Reproduction. Reprod. Sci. 29, 849–856 (2022).

13. Guo, X.-Y. et al. Cardiovascular and metabolic profiles of offspring conceived by assisted reproductive technologies: a systematic review and meta-analysis. Fertil Steril 107, 622–631.e5 (2017).

14. Hart, R. J. & Wijs, L. A. The longer-term effects of IVF on offspring from childhood to adolescence. Front Reprod Health 4, 1045762 (2022).

15. Li, H. & Auwerx, J. Mouse systems genetics as a prelude to precision medicine. Trends Genet 36, 259–272 (2020).

16. Rhon-Calderon, E. A. et al. Trophectoderm biopsy of blastocysts following IVF and embryo culture increases epigenetic dysregulation in a mouse model. Human Reproduction dead238 (2023) doi:10.1093/humrep/dead238.

17. Cui, L. et al. Increased risk of metabolic dysfunction in children conceived by assisted reproductive technology. Diabetologia 63, 2150–2157 (2020).

18. Kianpour, M. et al. Metabolic Syndrome and Assisted Reproductive Techniques. J Family Reprod Health 17, 80–85 (2023).

19. Qin, N. et al. Abnormal Glucose Metabolism in Male Mice Offspring Conceived by in vitro Fertilization and Frozen-Thawed Embryo Transfer. Front. Cell Dev. Biol. 9, (2021).

20. Heber, M. F. & Ptak, G. E. The effects of assisted reproduction technologies on metabolic health and disease†. Biol Reprod 104, 734–744 (2020).

21. Aljahdali, A. et al. The duration of embryo culture after mouse IVF differentially affects cardiovascular and metabolic health in male offspring. Human Reproduction 35, 2497–2514 (2020).

22. Rhon-Calderon, E. A. et al. In Vitro Fertilization induces reproductive changes in male mouse offspring and has multigenerational effects. 2024.11.06.622317 Preprint at 10.1101/2024.11.06.622317 (2024).

23. Zhou, X., Sun, Y., Feng, W., Wan, W. & Cui, L. Long-term effects of IVF on offspring kidneys in mice: observations from adolescence to adulthood. Reproductive BioMedicine Online 51, 104501 (2025).

24. Rhon-Calderon, E. A. et al. In Vitro Fertilization induces reproductive changes in male mouse offspring and has multigenerational effects. https://insight.jci.org/articles/view/188931/pdf (2025) doi:10.1172/jci.insight.188931.

25. Geraci, A. et al. Sarcopenia and Menopause: The Role of Estradiol. Front. Endocrinol. 12, (2021).

26. Emori, C. et al. Cooperative Effects of 17β-Estradiol and Oocyte-Derived Paracrine Factors on the Transcriptome of Mouse Cumulus Cells. Endocrinology 154, 4859–4872 (2013).

27. Liu, T. et al. Mouse model of menstruation: An indispensable tool to investigate the mechanisms of menstruation and gynaecological diseases (Review). Molecular Medicine Reports 22, 4463–4474 (2020).

28. Chiang, J. L. et al. Mitochondria in Ovarian Aging and Reproductive Longevity. Ageing Res Rev 63, 101168 (2020).

29. Robker, R. L., Hennebold, J. D. & Russell, D. L. Coordination of Ovulation and Oocyte Maturation: A Good Egg at the Right Time. Endocrinology 159, 3209–3218 (2018).

30. Cui, J. & Wang, Y. Premature ovarian insufficiency: a review on the role of tobacco smoke, its clinical harm, and treatment. Journal of Ovarian Research 17, 8 (2024).

31. Soygur, B. et al. A Roadmap for Three-Dimensional Analysis of the Intact Mouse Ovary. Methods Mol Biol 2677, 203–219 (2023).

32. Landry, D. A., Vaishnav, H. T. & Vanderhyden, B. C. The significance of ovarian fibrosis. Oncotarget 11, 4366–4370 (2020).

33. Amargant, F. et al. Ovarian stiffness increases with age in the mammalian ovary and depends on collagen and hyaluronan matrices. Aging Cell 19, e13259 (2020).

34. Shi, Y.-Q. et al. Premature ovarian insufficiency: a review on the role of oxidative stress and the application of antioxidants. Front Endocrinol (Lausanne) 14, 1172481 (2023).

35. Mu, L. et al. Physiological premature aging of ovarian blood vessels leads to decline in fertility in middle-aged mice. Nat Commun 16, 72 (2025).

36. Tharp, M. E., Malki, S. & Bortvin, A. Maximizing the ovarian reserve in mice by evading LINE-1 genotoxicity. Nat Commun 11, 330 (2020).

37. Wang, K. et al. Epigenetic regulation of aging: implications for interventions of aging and diseases. Sig Transduct Target Ther 7, 374 (2022).

38. Liu, Y. & Gao, J. Reproductive aging: biological pathways and potential interventive strategies. Journal of Genetics and Genomics 50, 141–150 (2023).

39. Nie, L., Wang, X., Wang, S., Hong, Z. & Wang, M. Genetic insights into the complexity of premature ovarian insufficiency. Reprod Biol Endocrinol 22, 94 (2024).

40. Kiriakidou, M., McAllister, J. M., Sugawara, T. & Strauss, J. F. Expression of steroidogenic acute regulatory protein (StAR) in the human ovary. J Clin Endocrinol Metab 81, 4122– 4128 (1996).

41. Schröder, S. K., Krizanac, M., Kim, P., Kessel, J. C. & Weiskirchen, R. Ovaries of estrogen receptor 1-deficient mice show iron overload and signs of aging. Front Endocrinol (Lausanne) 15, 1325386 (2024).

42. Gong, Y. et al. Age-related decline in the expression of GDF9 and BMP15 genes in follicle fluid and granulosa cells derived from poor ovarian responders. J Ovarian Res 14, 1 (2021).

43. López-Otín, C., Blasco, M. A., Partridge, L., Serrano, M. & Kroemer, G. Hallmarks of aging: An expanding universe. Cell 186, 243–278 (2023).

44. Rubina, K. et al. T-Cadherin (CDH13) and Non-Coding RNAs: The Crosstalk Between Health and Disease. International Journal of Molecular Sciences 26, 6127 (2025).

45. Pansera, M., Neeraj, N., Siddappa, D., Schuermann, Y. & Duggavathi, R. Expression of the glucose transporter 1 is associated with increased glucose uptake by granulosa cells during ovulation in mice. Theriogenology 236, 13–20 (2025).

46. Trigg, N. A. & Conine, C. C. Epididymal acquired sperm microRNAs modify post-fertilization embryonic gene expression. Cell Reports 43, 114698 (2024).

47. Orvieto, R. et al. Impact of aging on gene expression in human oocytes: a comparative analysis of young and older patients. Reprod Biol Endocrinol 23, 111 (2025).

48. Jiapaer, Z. et al. Regulation and roles of RNA modifications in aging related diseases. Aging Cell 21, e13657 (2022).

49. Bao, S., Yin, T. & Liu, S. Ovarian aging: energy metabolism of oocytes. Journal of Ovarian Research 17, 118 (2024).

50. Zhang, Y. et al. KIF20A Regulates Porcine Oocyte Maturation and Early Embryo Development. PLOS ONE 9, e102898 (2014).

51. Neyroud, A. S. et al. LARS2 variants can present as premature ovarian insufficiency in the absence of overt hearing loss. Eur J Hum Genet 31, 453–460 (2023).

52. Mehlmann, L. M. et al. The Gs-linked receptor GPR3 maintains meiotic arrest in mammalian oocytes. Science 306, 1947–1950 (2004).

53. Pyun, J.-A., Cha, D. H. & Kwack, K. LAMC1 gene is associated with premature ovarian failure. Maturitas 71, 402–406 (2012).

54. Rah, H. et al. Association of polymorphisms in microRNA machinery genes (DROSHA, DICER1, RAN, and XPO5) with risk of idiopathic primary ovarian insufficiency in Korean women. Menopause 20, 1067–1073 (2013).

55. Harris, S. E. et al. INHA promoter polymorphisms are associated with premature ovarian failure. Mol Hum Reprod 11, 779–784 (2005).

56. Bai, L. et al. ALKBH5 controls the meiosis-coupled mRNA clearance in oocytes by removing the N 6-methyladenosine methylation. Nat Commun 14, 6532 (2023).

57. Sutherland, J. M. et al. Knockout of RNA Binding Protein MSI2 Impairs Follicle Development in the Mouse Ovary: Characterization of MSI1 and MSI2 during Folliculogenesis. Biomolecules 5, 1228–1244 (2015).

58. Xu, J. & Gridley, T. Notch2 is required in somatic cells for breakdown of ovarian germ-cell nests and formation of primordial follicles. BMC Biology 11, 13 (2013).

59. Kim, H.-R. et al. Estrogen induces EGR1 to fine-tune its actions on uterine epithelium by controlling PR signaling for successful embryo implantation. The FASEB Journal 32, 1184– 1195 (2018).

60. Lu, S. et al. Repetitive Element DNA Methylation is Associated with Menopausal Age. Aging and disease 9, 435–443 (2018).

61. Gardini, E. S. et al. Differential ESR1 Promoter Methylation in the Peripheral Blood— Findings from the Women 40+ Healthy Aging Study. Int J Mol Sci 21, 3654 (2020).

62. Wheeler, A. C. et al. Associated features in females with an FMR1 premutation. Journal of Neurodevelopmental Disorders 6, 30 (2014).

63. Harner, R. et al. Ovulation induction is associated with altered growth but with preservation of normal metabolic function in murine offspring. F S Sci 2, 259–267 (2021).

64. Broekmans, F. J., Soules, M. R. & Fauser, B. C. Ovarian Aging: Mechanisms and Clinical Consequences. Endocrine Reviews 30, 465–493 (2009).

65. Salameh, Y., Bejaoui, Y. & El Hajj, N. DNA Methylation Biomarkers in Aging and Age-Related Diseases. Front. Genet. 11, (2020).

66. Wang, K. et al. Epigenetic regulation of aging: implications for interventions of aging and diseases. Sig Transduct Target Ther 7, 1–22 (2022).

67. Campen, K. A., McNatty, K. P. & Pitman, J. L. A protective role of cumulus cells after short-term exposure of rat cumulus cell-oocyte complexes to lifestyle or environmental contaminants. Reproductive Toxicology 69, 19–33 (2017).

68. Gonnella, F. et al. A Systematic Review of the Effects of High-Fat Diet Exposure on Oocyte and Follicular Quality: A Molecular Point of View. Int J Mol Sci 23, 8890 (2022).

69. Al-Yasari, A., Jabbar, S., Cabrera, M. A., Rousseau, B. & Sarkar, D. K. Preconception Alcohol Exposure Increases the Susceptibility to Diabetes in the Offspring. Endocrinology 162, bqaa188 (2020).

70. Xu, X., Pang, Y. & Fan, X. Mitochondria in oxidative stress, inflammation and aging: from mechanisms to therapeutic advances. Signal Transduct Target Ther 10, 190 (2025).

71. Behringer, R. Manipulating the Mouse Embryo: A Laboratory Manual. (Cold Spring Harbor Laboratory Press, Cold Spring Harbor, New York, 2014).

72. Myers, M., Britt, K. L., Wreford, N. G. M., Ebling, F. J. P. & Kerr, J. B. Methods for quantifying follicular numbers within the mouse ovary. 10.1530/rep.1.00095(2004) doi:10.1530/rep.1.00095.

73. Vrooman, L. A. et al. Assisted reproductive technologies induce temporally specific placental defects and the preeclampsia risk marker sFLT1 in mouse. Development 147, (2020).

74. Prasasya, R. D. et al. Iterative oxidation by TET1 is required for reprogramming of imprinting control regions and patterning of mouse sperm hypomethylated regions. Developmental Cell 59, 1010–1027.e8 (2024).

75. Quiros, P. M., Goyal, A., Jha, P. & Auwerx, J. Analysis of mtDNA/nDNA ratio in mice. Curr Protoc Mouse Biol 7, 47–54 (2017).

76. Zhang, B. A., et al. Comprehensive Transcriptomic and Epigenomic Insights into Environmental Toxicant Exposures: The TaRGET II Resource. bioRxiv 2025.07.28.667191 (2025) doi:10.1101/2025.07.28.667191.

