## Extended Data Fig. for "In Vitro Fertilization Accelerates Female Reproductive Aging Through Early Ovarian Failure"

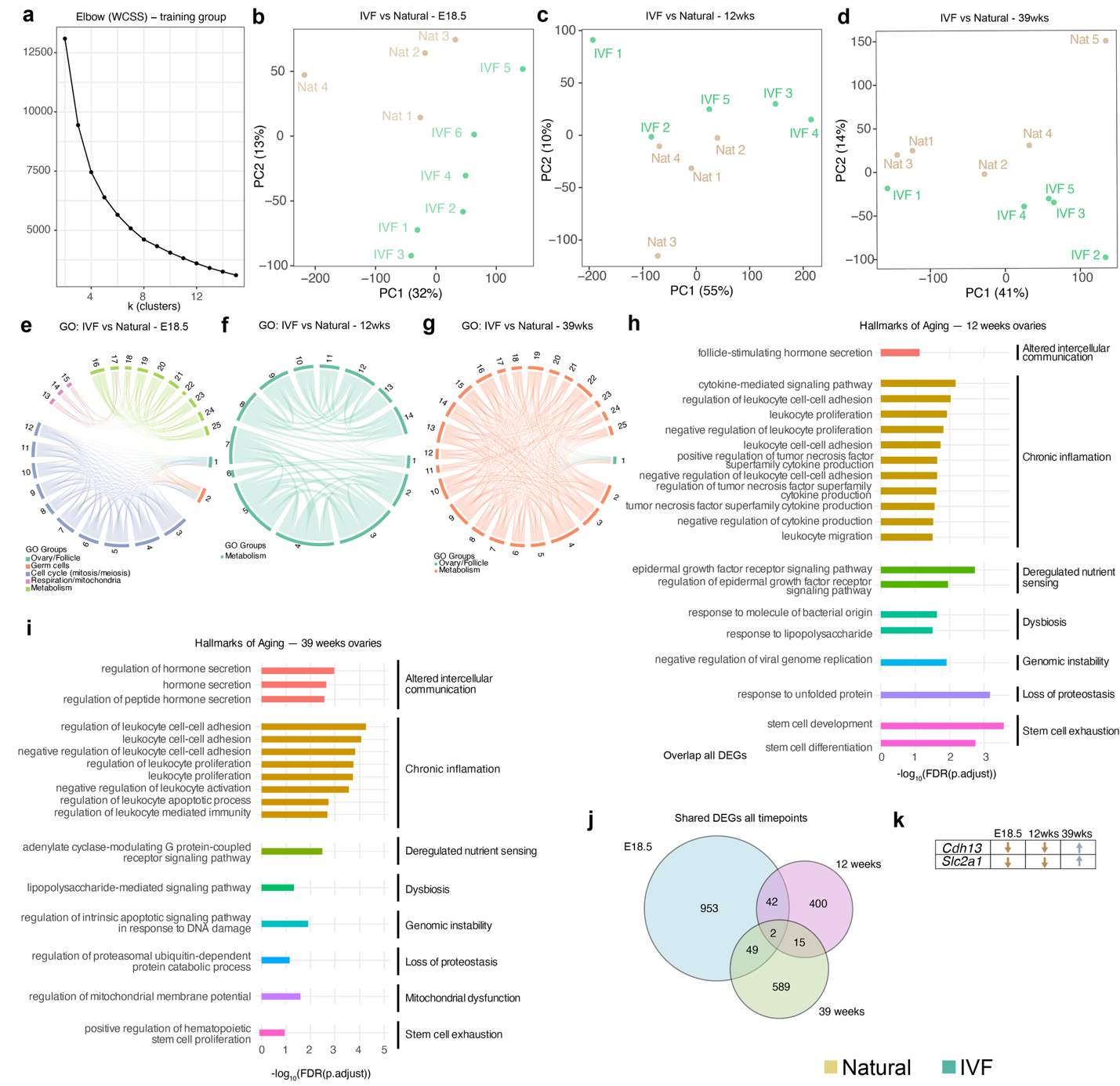

**Extended Data 1. Corresponding to main Figure 3.**

**a,** Elbow: plot analysis for k selection in WCSS analysis. Principal Component Analysis (PCA) for whole ovary RNA-seq in Figure 3: **b,** E18.5; **c,** 12-weeks; **d,** 39-weeks. Chord plots for Network representation of most affected GO terms at **e,** E18.5; **f,** 12-weeks; **g,** 39-weeks. **h,** Venn diagram to find the overlap genes at the three developmental timepoints. **i,** List of genes that are shared between the different timepoints and the direction of their dysregulation.

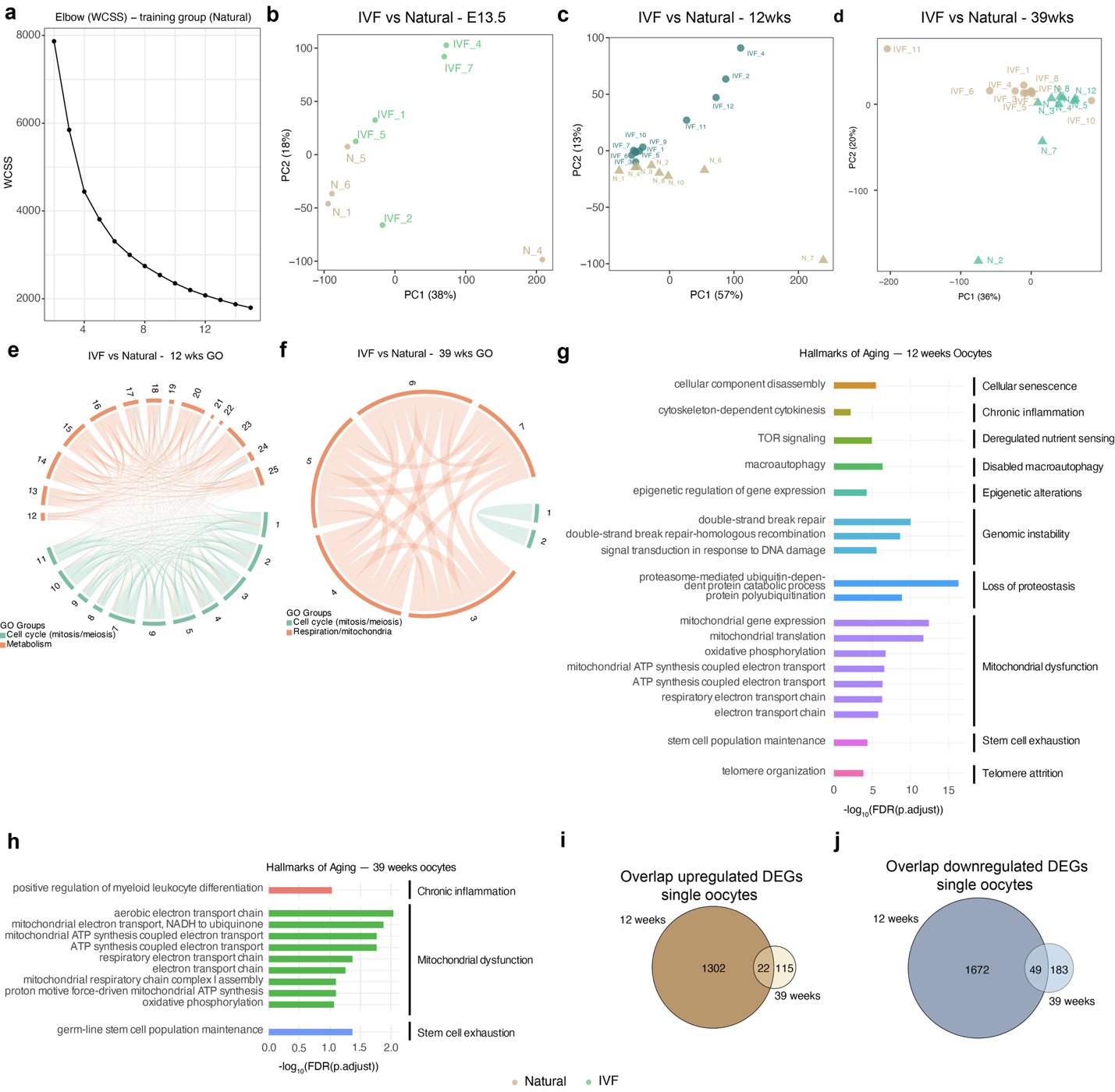

**Extended Data 2. Corresponding to main Figure 4.**

**a,** Elbow: plot analysis for k selection in WCSS analysis. Principal Component Analysis (PCA) for primordial germ cells and oocytes RNA-seq in Figure 4: **b,** E13.5; **c,** 12-weeks; **d,** 39-weeks. Chord plots for Network representation of most affected GO terms at **e,** E13.5; **f,** 12-weeks; **g,** 39-weeks. Overlap between 12 and 39 weeks for **g,** upregulated DEGs and **g,** downregulated DEGs.

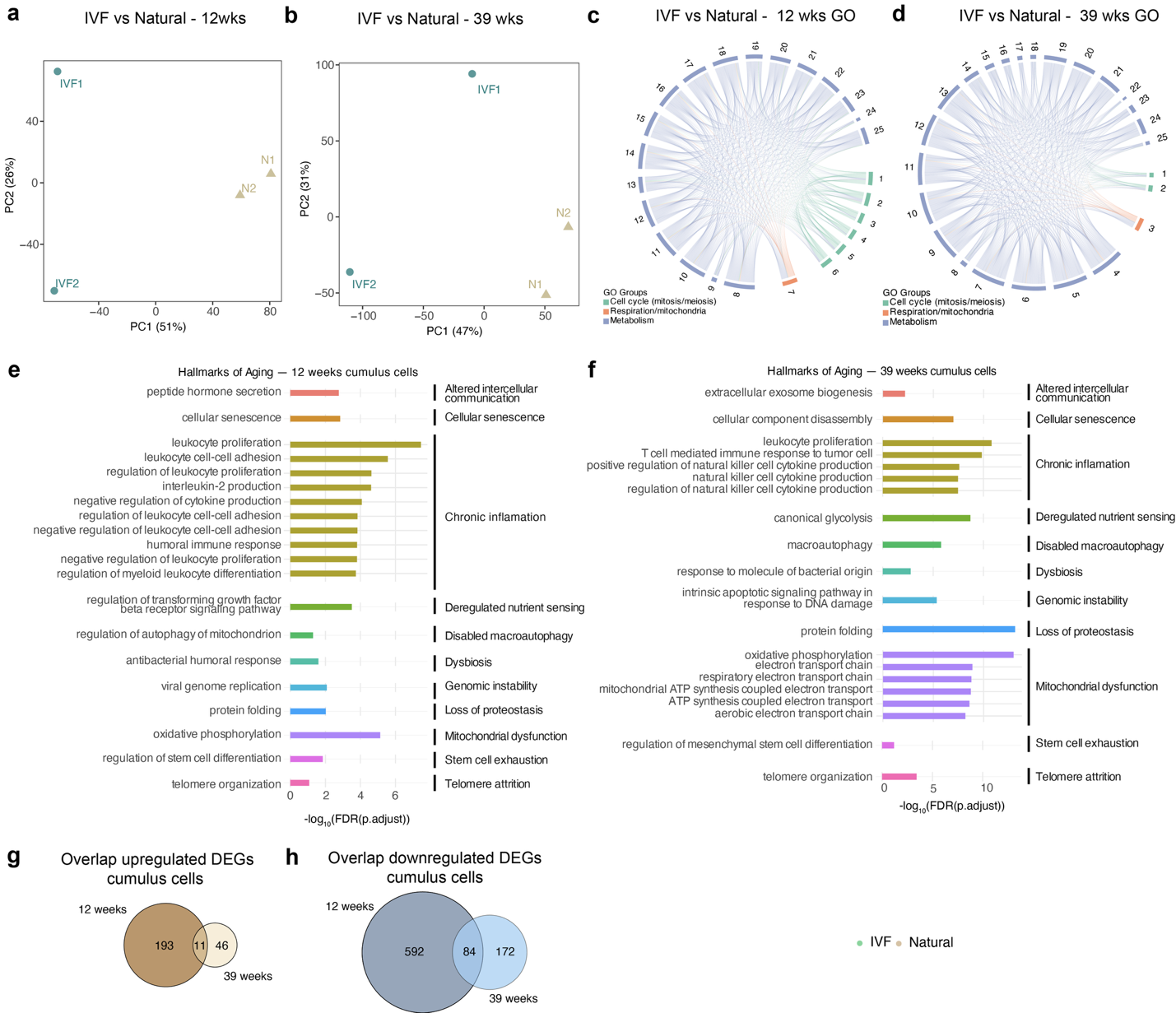

**Extended Data 3. Corresponding to main Figure 5.**

Principal Component Analysis (PCA) for cumulus cells RNA-seq in Figure 4: **a,** 12 weeks; **b,** 39 weeks. Chord plots for Network representation of most affected GO terms at **c,** 12 weeks; **d,** 39 weeks. Gene ontology to identify hallmark of aging pathways based on López-Otín et al.2023: **e,** 12 weeks and **f,** 39 weeks. Overlap between 12 and 39 weeks for **g,** upregulated DEGs and **h,** downregulated DEGs.
